# “Comparative Analysis of Gut Microbiome in Individuals of the Old Civilization of Caral-Supe Based on Data from 16S rRNA and ITS region”

**DOI:** 10.1101/2020.04.24.060582

**Authors:** Andrés Vásquez-Domínguez, Luis Jaramillo-Valverde, Kelly S. Levano, Pedro Novoa-Bellota, Marco Machaguay-Romero, Ruth Garcia-de-la-Guarda, Raul J. Cano, Ruth Shady Solis, Heinner Guio

## Abstract

Genetic and microbiome studies of ancient Caral-Supe civilization have not yet been published. For this reason, the objective of this work is to identify the microorganisms and possible diseases that existed in this ancient civilization using coprolites samples. To do this, two coprolites samples were analyzing through high-throughput sequencing data of 16S rRNA gene and an intergenic region (ITS).

## CONTENT

The Caral-Supe civilization (3500 BC), the oldest in Peru and America and contemporary to the Mesopotamia, Egypt, India and China civilizations (1), was discovered by anthropologist and archeologist Dr. Ruth Shady.

The Caral-Supe population relied both in agricultural and marine products, permanently available through a local exchange network (2). Genetic and microbiome studies of the Caral civilization have not yet been published. In the present study, we characterized the gut microbiome of this civilization using coprolites samples found near the Sacred City of Caral. Coprolites (fossilized fecal material) provide information on the composition of the intestinal microbiota, and thus help us obtain information of cultural practices, dietary habits, and individual’s gut microbiota state (3). High-throughput sequence analysis of the 16S rRNA gene and an intergenic region (ITS) was performed to study two coprolites samples (PACS-176 and PEACS-329), provided by the Zona Arqueológica Caral. Molecular processes were carried out at the INBIOMEDIC laboratory. Biosecurity measures and standard precautions were used for ancient DNA work. Prior to DNA extraction, samples were prepared by discarding the outermost portions of the coprolite samples to eliminate risk of contamination, and a replica of each sample was obtained for further analysis. Ancient genomic DNA was extracted using the PowerSoil DNA Isolation Kit from Qiagen, following the manufacturer’s instructions (4).

The V4 variable region of bacterial 16S rRNA genes was targeted for high throughput sequencing using predetermined primers 515F (GTGYCAGCMGCCGCGGTAA) and 806R (GGACTACNVGGGTWTCTAAT). In addition, for genetic identification of fungi, ITS analysis was performed using primers ITS1 (CTTGGTCATTTAGAGGAAGTAA) and ITS4 (GCTGCGTTCTTCATCGATGC) (5).

Library construction and sequencing was performed in MR-DNA laboratory (www.mrdnalab.com, Shallowater, TX, USA), and sequences were determined using an Illumina MiSeq instrument following the manufacturer’s guidelines. Barcodes, sequences <150 bp, sequences with ambiguous bases, and homopolymer runs greater than 6 bp were removed; using the open access software www.mrdnafreesoftware.com. Operative taxonomic units (OTU) were generated and chimeras were filtered. The OTUs were defined by grouping at a 3% divergence (97% similarity). The final OTUs were taxonomically classified using BLASTn against a database derived from RDPII (http://rdp.cme.msu.edu) and NCBI (www.ncbi.nlm.nih.gov).

In both coprolites samples examined, the most representative OTUs at the phylum level for 16S rRNA bacterial genes was the Firmicutes (PEACS-329 with 36.44% and PACS-176 with 98.91%) (Fig 1). Similarly, for ITS analysis, the most prevalent phylum was Ascomycota (PEACS-329 with 99.92% and PACS-176 with 99.99%) (Fig 2). Firmicutes is one of the most frequent phylum in the human intestinal microbiota and is related to cardiovascular problems (6).

**FIG 1.**
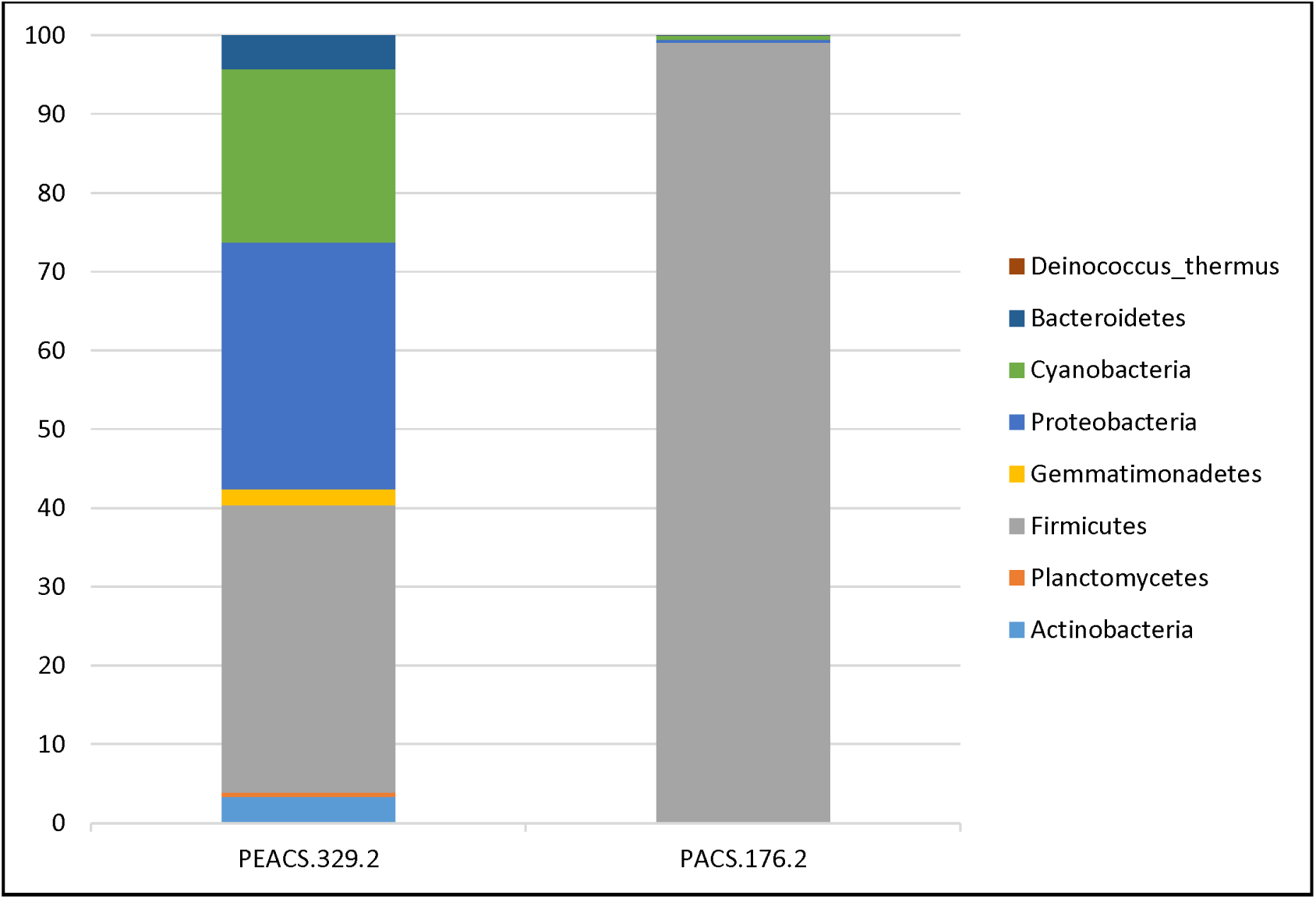
Bar chart representing bacterial diversity of coprolites based on 16S rRNA amplicon analysis. The relative abundance (%) at the phylum level is shown.

**FIG 2.**
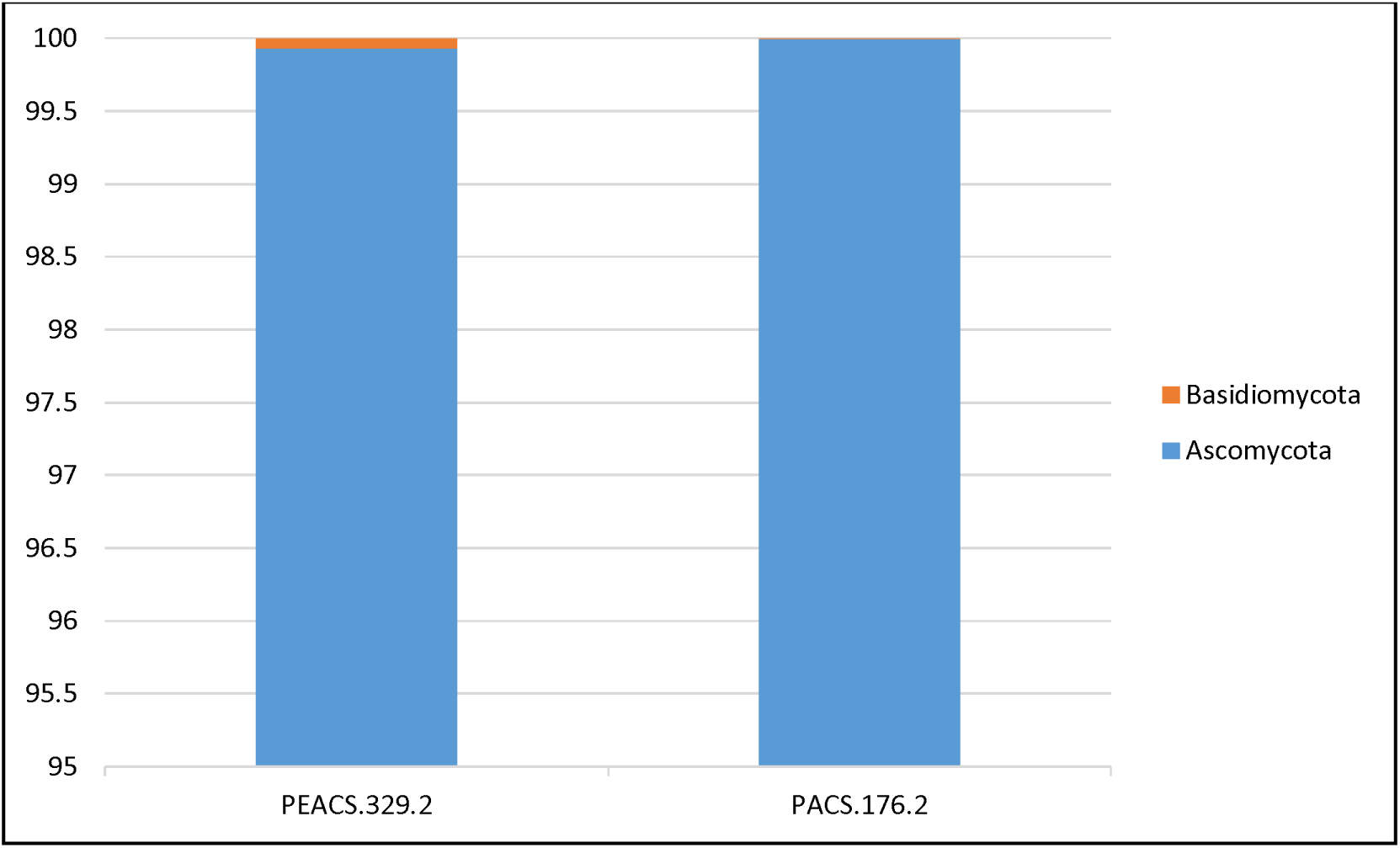
Bar chart representing fungal diversity of coprolites based on ITS amplicon analysis. The relative abundance (%) at the phylum level is shown.

At the genus level for 16S rRNA bacterial genes, the OTUs showed *Halospirulina sp.* to be the most representative (21.96%) for the sample PEACS-329 and *Virgibacillus sp.* with 87.04% for the PACS-176 sample (Table 1). For ITS analysis, the most frequent OTU was *Phialosimplex sp.* (PEACS-329 with 77.49% and PACS-176 with 74.88%) (Table 2). *Virgibacillus sp.* is a well representative genus of the phylum Firmicutes (7), which is the phylum most frequently found in both samples.

**TABLE 1:**
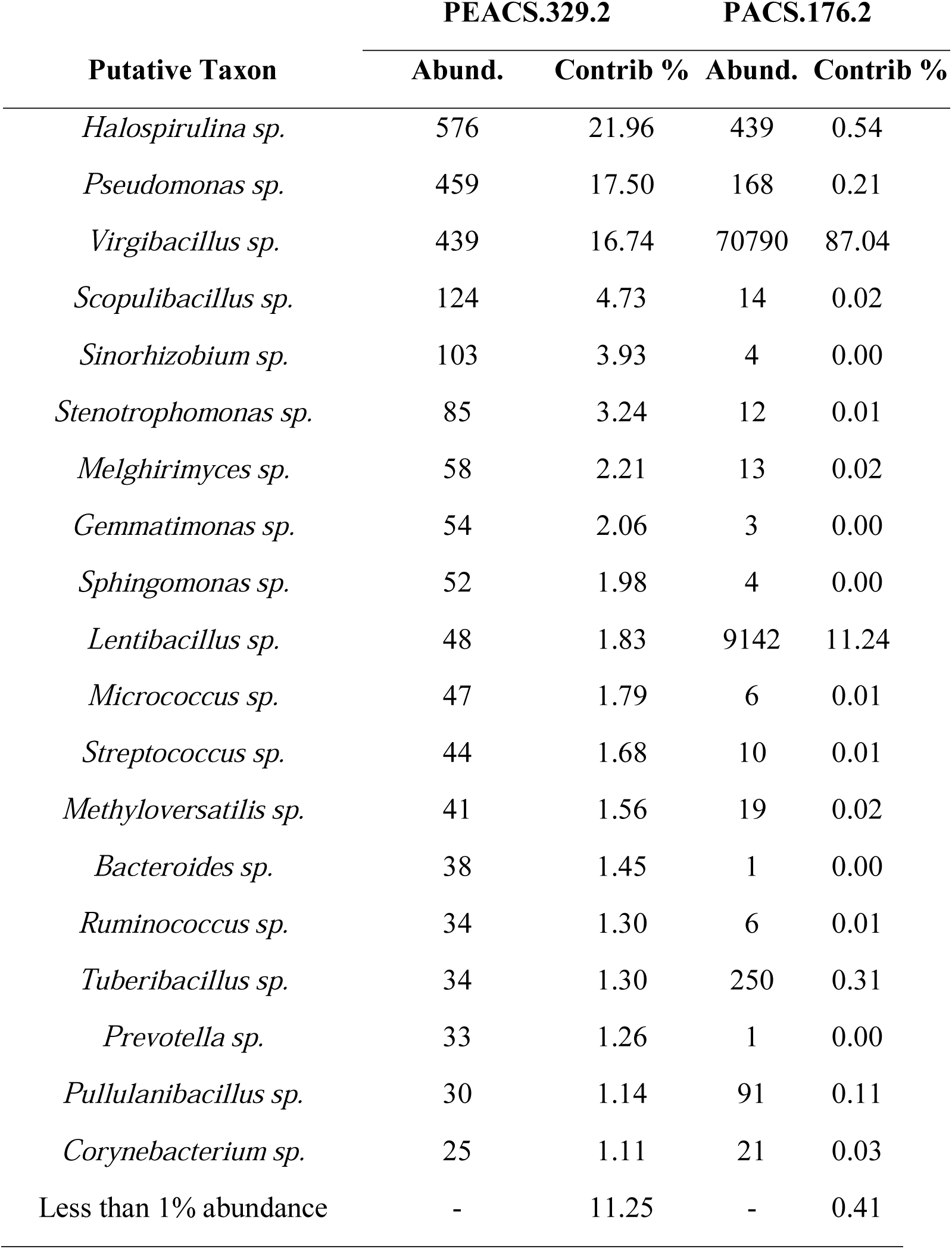
Analysis of Bacterial taxa impacting clustering of PEACS 329.2 and PACS 176.2 Coprolite.

**TABLE 2:** Analysis of Fungal taxa impacting clustering of PEACS 329.2 and PACS 176.2 Coprolites.

| Putative Taxon | PEACS.329.2 |  | PACS.176.2 |  |
| --- | --- | --- | --- | --- |
|  | Abund. | Contrib % | Abund. | Contrib % |
| *Phialosimplex sp.* | 89689 | 77.49 | 48746 | 74.88 |
| *Aspergillus sp.* | 17775 | 15.36 | 15262 | 23.44 |
| *Epicoccum sp.* | 7752 | 6.70 | 306 | 0.47 |
| *Penicillium sp.* | 245 | 0.21 | 11 | 0.02 |
| *Saccharomyces sp.* | 124 | 0.11 | 8 | 0.01 |
| *Malassezia sp.* | 82 | 0.07 | 5 | 0.01 |
| *Fusarium sp.* | 67 | 0.06 | 751 | 1.15 |
| *Phoma sp.* | 8 | 0.01 | 0 | 0.00 |
| *Alternaria sp.* | 0 | 0.00 | 11 | 0.02 |

The present study has allowed the development of a comparative analysis of gut microbial diversity in coprolites from the Caral civilization. Further analysis will be carried out with additional coprolites samples to further identify organisms that colonized the gut microbiome of the Caral-Supe ancient inhabitants; and in this way be able to determine the lifestyle (diet) and its influence on the gut microbial communities.

## ACKNOWLEDGMENTS

The Fondo Nacional de Desarrollo Científico, Tecnológico y de Innovación Tecnológica (FONDECYT) and BANCO MUNDIAL with contract number E043-2018-FONDECYT/BM-IADT-AV., supported this study The funders had no role in the study design, the data collection and interpretation, or the decision to submit the work for publication.

